# Calculation of fungal and bacterial inorganic nitrogen immobilization rates in soil

**DOI:** 10.1101/2020.03.19.996876

**Authors:** Xiaobo Li, Hongbo He, Xudong Zhang, Caner Kazanci, Zhian Li, Magdalena Necpalova, Qianqian Ma

**Author notes:** Corresponding author: Q, Ma, South China Botanical Garden, Chinese Academy of Sciences, Guangzhou 510650, China. *E-mail.

## Abstract

Microbial inorganic nitrogen (N) immobilization is an important mechanism in the retention of N in soils. However, as a result of the high diversity and complexity of soil microorganisms, there is still no effective approach to measuring the respective immobilization rates of inorganic N by fungi and bacteria, which are the two dominant microbial communities in soils. We propose a mathematical framework, combining the experimentally measurable gross inorganic N immobilization rate and proxies for fungal and bacterial inorganic N immobilization rates, to quantify the respective immobilization rates of inorganic N by fungal and bacterial communities in soil. Our approach will help to unravel the mechanisms of N retention in soils.

---

The microbial immobilization of inorganic nitrogen (N) has a vital role in controlling the size of the soil inorganic N pool and is therefore an important mechanism for the retention of N in ecosystems (Davidson et al 1992, Stark and Hart 1997, Zhang et al 2013, Zogg et al 2000). Through this immobilization process, inorganic N in soil is converted to microbial biomass N and subsequently re-mineralized or converted to stable organic N, eventually reducing the risk of N losses from soil (Recous et al 1990, Tahovská et al 2013, Zhang et al 2019). As the dominant microorganisms in soil, fungi and bacteria are probably the main participants in inorganic N immobilization (Bottomley et al 2012, Boyle et al 2008, Myrold and Posavatz 2007). Given the distinct physiologies, morphologies, lifestyles and quantities of these two microbial groups in soil (Lauber et al 2008, Rousk and Bååth 2011, Six et al 2006, Waring et al 2013), the relative importance of fungi and bacteria in soil inorganic N immobilization is likely to be unequal (Bottomley et al 2012, Li et al 2019, Myrold and Posavatz 2007). However, as a result of the high diversity and complexity of soil microorganisms, quantifying the respective rates of immobilization of inorganic N by fungal and bacterial communities in soil is challenging (Fierer 2017, Li et al 2019, Li et al 2020), although the gross inorganic N immobilization rate can be measured using well-established ^15^N isotope techniques (e.g., the ^15^N pool dilution method) (Cheng et al 2017, Murphy et al 2003).

Amino sugars, which are important constituents of microbial cell walls, have different origins in microorganisms. Among the amino sugars identified in microorganisms, muramic acid (MurN) originates exclusively from bacterial peptidoglycan, whereas glucosamine (GlcN) is mainly in the form of chitin in fungal cell walls (Amelung 2001, Parsons 1981, Zhang and Amelung 1996). Based on their microbial source specificity, stable isotope probing based on amino sugars (^15^N-AS-SIP) has been developed to disentangle the immobilization processes of inorganic N by fungi and bacteria in soils (He et al 2006, He et al 2011a, He et al 2011b, Liang and Balser 2010, Reay et al 2019a, Reay et al 2019b).

This approach has now been extended to indicate the inorganic N immobilization rates of fungal and bacterial communities in soils (Li et al 2019, Li et al 2020). More specifically, given the relatively long persistence of amino sugars in soils (mean turnover time >2 years, much longer than that of the living microorganisms) (Derrien and Amelung 2011, Glaser et al 2006, Liu et al 2016), the newly formed ^15^N-labeled amino sugars are considered to be stable in soil even after cell death (Glaser et al 2004, Gunina et al 2017). The fungal-derived ^15^N-GlcN and bacterial-derived ^15^N-MurN synthesis rates within a short period of incubation after ^15^N tracer addition have therefore been used as proxies for the rates of immobilization of inorganic N by fungi and bacteria, respectively (Li et al 2019, Li et al 2020). However, mainly as a result of the variation in the composition of tissues of massive microbial species, but also within each species under different growth conditions, the actual contents of GlcN and MurN in the respective biomasses of fungi and bacteria in soil are almost unobtainable (Appuhn and Joergensen 2006, Engelking et al 2007, Glaser et al 2004, Joergensen 2018). It is also still unclear how fast do the cell N-containing components turn over intracellularly and extracellularly in soil (Dippold et al 2019, Engelking et al 2007, Gunina et al 2017). As a consequence, converting the synthesis rates of ^15^N-labeled amino sugars specific for fungi and bacteria to the actual inorganic N immobilization rates in soil is challenging.

To bypass this intractable problem, we propose a mathematical framework to estimate the conversion coefficients between fungal and bacterial inorganic N immobilization rates and their respective proxies by combining the gross inorganic N immobilization rate with proxies for the respective inorganic N immobilization rates of fungi and bacteria. In this way, we can obtain the respective immobilization rates of inorganic N by fungal and bacterial communities in soil.

## Calculation of fungal and bacterial inorganic N immobilization rates

Our proposed calculation is based on the assumption that fungi and bacteria are the dominant participants in soil microbial inorganic N immobilization. If both the gross inorganic N immobilization rate and the proxies for inorganic N immobilization rates of fungi and bacteria have been measured on *n* soil samples (*n*≥2), then the respective immobilization rates of inorganic N by fungal and bacterial communities can be calculated.

The measured variables are:

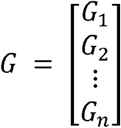: gross microbial inorganic N immobilization rates for *n* samples (mg N kg^−1^ day^−1^);

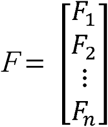: fungal-derived ^15^N-GlcN synthesis rates for *n* samples (mg N kg^−1^ day^−1^);

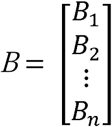: bacterial-derived ^15^N-MurN synthesis rates for *n* samples (mg N kg^−1^ day^−1^).

The two parameters to be estimated are:

*K*_*F*_: the conversion coefficient from the fungal-derived ^15^N-GlcN synthesis rate to the fungal inorganic N immobilization rate;

*K*_*B*_: the conversion coefficient from the bacterial-derived ^15^N-MurN synthesis rate to the bacterial inorganic N immobilization rate.

Using the ^15^N-labeled amino sugars synthesis rates and conversion coefficients, the estimated fungal and bacterial inorganic N immobilization rates (mg N kg^−1^ day^−1^) are, respectively, calculated as:

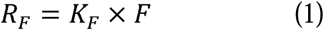

And

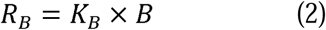

Their sum is therefore the estimated gross microbial inorganic N immobilization rate (mg N kg^−1^ day^−1^):

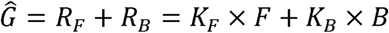

The measured gross microbial inorganic N immobilization rate results are included in the equation:

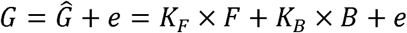

where *e* is the estimation error. This equation can be rewritten in a matrix format:

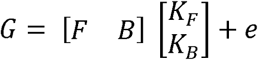

Alternatively,

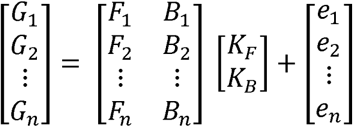

If we let 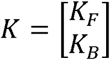 and 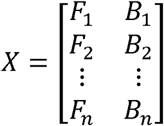, we obtain:

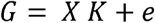

The least-squares estimators that minimize the sum of the squared residuals are given in the following (see Appendix for the detailed derivation) (Wackerly et al 2014):

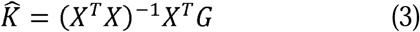

To illustrate how this approach works, we calculated the soil nitrate (NO_3_ ^−^) immobilization rates of fungi and bacteria using the gross NO_3_^−^ immobilization rates reported by Zhang et al (2013) and the ^15^N-labeled amino sugars synthesis rates reported by Li et al (2019). Both studies studied the effect of land conversion from forest to agriculture on the soil NO_3_^−^ immobilization in subtropical zones of China. Ideally, the gross NO_3_^−^ immobilization rates and the ^15^N-labeled amino sugars synthesis rates should be measured under the same experimental conditions such as sampling sites. Due to the unavailability of such data, we roughly treat selected studies as being conducted at the same sites. Therefore, the results in Table 1 are presented as an illustrative example, rather than as reliable estimates. For simplicity, only the mean rates for forest and agricultural lands were used in this example (*n* = 2). The conversion coefficients were obtained by substituting the measured gross NO_3_^−^ immobilization rates and the ^15^N-labeled amino sugars synthesis rates into Equation (3). The fungal and bacterial NO_3_^−^ immobilization rates were then calculated using Equations (1) and (2). A summary of measured data and estimated values is provided in Table 1.

**Table 1.** An illustration of the method of calculating soil fungal and bacterial NO_3_^−^ immobilization rates under different land use scenarios. The gross NO_3_^−^ immobilization rates (*G*) were obtained from Zhang et al (2013). The synthesis rates of fungal-derived ^15^N-GlcN (*F*) and bacterial-derived ^15^N-MurN (*B*) were obtained from Li et al (2019) (see Table S1 for detailed calculations). These values are presented as an illustrative example, rather than as reliable estimates.

| Land use | $G$ | $F$ | $B$ | $K_F$ | $K_B$ | $R_F$ | $R_B$ |
| --- | --- | --- | --- | --- | --- | --- | --- |
| | $\text{mg N kg}^{-1} \text{ day}^{-1}$ | | | | | $\text{mg N kg}^{-1} \text{ day}^{-1}$ | |
| Woodland | 0.47 | 0.0303 | 0.0022 | 13.78 | 23.83 | 0.42 | 0.05 |
| Agriculture | 0.10 | 0.0057 | 0.0009 | 13.78 | 23.83 | 0.08 | 0.02 |

Note: $K_F$ and $K_B$ are the conversion coefficients between $F$ , $B$ and the $\text{NO}_3^-$ immobilization rates of fungi ( $R_F$ ) and bacteria ( $R_B$ ), respectively.

The results showed that the NO_3_^−^ immobilization rates of fungi in woodland and agricultural soils were about 8.4 and four times those of bacteria, indicating that fungi dominated the microbial NO_3_ ^−^ immobilization in the studied soil (Table 1). Conversion to agricultural use led to decreases in the fungal and bacterial NO_3_^−^ immobilization rates of 0.34 and 0.03 mg N kg^−1^ day^−1^, respectively, which suggests that the decrease in the fungal NO_3_^−^ immobilization rate dominates the decrease in the gross soil microbial NO_3_ ^−^ immobilization caused by the land use change.

## Advantages and limitations of this approach

Understanding the microbially mediated N cycling processes in soil is central to unraveling soil N retention mechanisms and has ramifications for reducing N losses and managing ecosystem productivity. As a result of the high diversity and complexity of microbial communities, quantifying the process rates of different microbial groups has been a great challenge, especially in soil (Bardgett and Van Der Putten 2014, Fierer 2017, Stres and Tiedje 2006). Our approach provides an effective way to mathematically, rather than mechanically, quantify the relative importance of fungal and bacterial communities in soil inorganic N immobilization. It circumvents the bottleneck of directly measuring or estimating the inorganic N immobilization rates of fungi and bacteria in soil. The experimentally accessible gross inorganic N immobilization rate and proxies of fungal and bacterial inorganic N immobilization rates are used to estimate the conversion coefficients between fungal and bacterial inorganic N immobilization rates and their respective proxies. The conversion coefficients obtained inherently take into account both the actual contents of GlcN and MurN in the respective biomasses of fungi and bacteria and the turnover of cell N-containing components in the studied soil. Because the rationale and mathematical derivation are universal, our method may also be applicable to other environmental systems, such as freshly colonized organic substrates (Appuhn and Joergensen 2006). This approach relies on the simplifying assumption that only fungi and bacteria are involved in soil microbial inorganic N immobilization. This assumption may not quite hold true, because Archaea may also contribute to inorganic N immobilization (Laughlin et al 2009). Considering that Archaea contain GlcN, but not MurN (Joergensen 2018), the contribution of Archaea, if any, is included in the fungal inorganic N immobilization rates by adopting our approach. Nevertheless, considering that Archaea account for less than <1% of the soil microbial biomass (Fierer 2017), the errors caused by this assumption are probably trivial.

## Conclusions

We propose a mathematical approach that combines the mechanically accessible gross inorganic N immobilization rate and proxies for fungal and bacterial inorganic N immobilization rates (measured by ^15^N-AS-SIP) to quantify the inorganic N immobilization rates of fungal and bacterial communities in soil. This approach, although not without its limitations, allows us for the first time to disentangle the actual contribution of fungi and bacteria to the immobilization of N-containing substrates in soil. Promisingly, integrating both fungal and bacterial inorganic N immobilization rates into terrestrial ecosystem models (e.g., microbial models) will improve our ability to understand, predict and manage the N retention capacity in soils under different scenarios (Waring et al 2013).

## Supporting information

Appendix: Derivation of least squares estimators

## Acknowledgments

This study was supported by the National Natural Science Foundation of China (41977097, 41401279, 31600392 and 41630862), the Natural Science Foundation of Guangdong Province (2019A1515012067 and 2016A030310013), R & D program of Guangdong Provincial Department of Science and Technology (2018B030324003), China Soil Microbiome Initiative of the Chinese Academy of Sciences (XDB15040200), the Research Fund of State Key Laboratory of Soil and Sustainable Agriculture, Nanjing Institute of Soil Science, Chinese Academy of Sciences (Y412201422), and National Key Research and Development Program of China (2017YFC0506305). H. He and X. Zhang acknowledge the CAS Interdisciplinary Innovation Team. X. Li appreciates China Scholarship Council for providing funds to support his study in Switzerland.

