## Appendix: Derivation of least squares estimators for "Calculation of fungal and bacterial inorganic nitrogen immobilization rates in soil"

*Corresponding author:

**Appendix: Derivation of least squares estimators**

$$G= X K+e$$

The vector of residual *e* is given as:

$$e=G-XK$$

The sum of squared residuals:

$$e^{T}e=\left( G-XK \right)^{T}\left( G-XK \right)$$

$$=G^{T}G-K^{T}X^{T}G-G^{T}XK+K^{T}X^{T}XK$$

Given that $K^{T}X^{T}G=\left( K^{T}X^{T}G \right)^{T}=G^{T}XK$. We get

$$e^{T}e=G^{T}G-2K^{T}X^{T}G+K^{T}X^{T}XK$$

To find the K that minimizes the sum of squared residuals, we need to take the derivative of $e^{T}e$ with respect to K:

$$\frac{{\partial e}^{T}e}{\partial K}=-2X^{T}G+2X^{T}XK$$

We set this to zero at the optimum, $\hat{K}$:

$$-2X^{T}G+2X^{T}X\hat{K}=0$$

We get the ‘normal equation’:

$$X^{T}X\hat{K}=X^{T}G$$

Solving this equation,

$${\left( X^{T}X \right)^{-1}X}^{T}X\hat{K}={\left( X^{T}X \right)^{-1}X}^{T}G$$

Thus, the least-squares estimates are:

$$\hat{K}={\left( X^{T}X \right)^{-1}X}^{T}G$$

**Table S1** Calculating the synthesis rates of fungal-derived ^15^N-GlcN (*F*) and bacterial-derived ^15^N-MurN (*B*) from Li et al (2019).

| Land use | F-^15^N-GlcN | ^15^N-MurN | Incubation Time | GlcN-N | MurN-N | *F* | *B* |
| --- | --- | --- | --- | --- | --- | --- | --- |
|  | mg kg^-1^ | | day | % | | mg N kg^-1^ d^-1^ | |
| Woodland | 7.75 | 0.79 | 20 | 7.82 | 5.58 | 0.0303 | 0.0022 |
| Agriculture | 1.47 | 0.31 | 20 | 7.82 | 5.58 | 0.0057 | 0.0009 |

**Note:** F-^15^N-GlcN and ^15^N-MurN are the accumulative amounts of newly formed Fungal-derived ^15^N-GlcN and bacterial-derived ^15^N-MurN after 20 days incubation, respectively. GlcN-N and MurN-N are N contents of GlcN and MurN (%), respectively.
